# Ectopically expressed *Airn* lncRNA deposits Polycomb with a potency that rivals *Xist*

**DOI:** 10.1101/2023.05.09.539960

**Authors:** Jackson B. Trotman, Aki K. Braceros, Steven R. Bischoff, McKenzie M. Murvin, Samuel P. Boyson, Rachel E. Cherney, Quinn E. Eberhard, Elizabeth W. Abrash, Dale O. Cowley, J. Mauro Calabrese

## Abstract

We report that when expressed at similar levels from an isogenic locus, the *Airn* lncRNA induces Polycomb deposition with a potency that rivals *Xist*. However, when subject to the same degree of promoter activation, *Xist* is more abundant and more potent than *Airn*. Our data definitively demonstrate that the *Airn* lncRNA is functional and suggest that *Xist* achieved extreme potency in part by evolving mechanisms to promote its own abundance.

## Main text

Long noncoding RNAs (lncRNAs) play critical roles in development by repressing transcription *in cis* (on the same chromosome from which they were transcribed ^1^). In the most extreme example, expression of the lncRNA *Xist* represses transcription across one entire X chromosome during the essential process of X inactivation. Critical for stable repression by *Xist* are enzymes called the Polycomb Repressive Complexes, PRC1 and PRC2, which monoubiquitylate lysine 119 of Histone H2A (H2AK119ub) and trimethylate lysine 27 of Histone H3 (H3K27me3), respectively ^2^. The mechanisms that *Xist* uses to silence gene expression and recruit PRCs to chromatin remain under investigation. However, there is little question that the *Xist* lncRNA product is functional; its ectopic expression from essentially any chromosomal location, not only the X, results in chromosome-wide transcriptional repression and PRC recruitment ^1^.

Beyond *Xist*, a handful of other *cis*-acting repressive lncRNAs have been identified. The most potent of these is a lncRNA called *Airn*, whose expression has been shown to repress genes and recruit PRCs within a genomic interval that spans roughly 15 megabases (Mb) in the extraembryonic tissues of the mouse ^3, 4^. *Airn* is an unusual RNA ^5^. Unlike *Xist*, which is robustly spliced, polyadenylated, and stable (half-life of ∼6 hours), *Airn* evades splicing, is incompletely polyadenylated, and is unstable (half-life of ∼1 hour; ^4, 6^). Also, while *Xist* is processed into distinct transcripts, *Airn* transcripts are variable in length, terminating gradually as distance from *Airn*’s transcription start site increases, upwards of 100kb away ^7^. These unusual properties raise the question of whether it is the *Airn* lncRNA product or act of *Airn* transcription that is responsible for its regulatory effect. This same question has been posed for other putative regulatory lncRNAs. In many cases, the act of transcription, and not the lncRNA product, has been found to be the dominant mediator of regulation ^8–11^.

To determine whether the *Airn* lncRNA product is functional, we sought to induce its expression from an ectopic locus; this would allow us to investigate effects arising only from expression of the lncRNA itself, decoupled from any effects of endogenous *cis*-regulatory features or chromosomal structures that could have evolved in concert with the *Airn* gene. To this end, we cloned the first ∼89kb of the *Airn* gene downstream of a doxycycline (dox)-inducible promoter in a DNA vector that enabled its Cre-mediated insertion into the C57BL/6J (B6) allele of the *Rosa26* locus on chromosome 6 (chr6), in F1-hybrid mouse embryonic stem cells (ESCs) derived from a cross between B6 and CAST/EiJ (CAST) mice ^12^ (Figure 1A). We obtained two separate ESC clones containing the *Airn* gene inserted into B6 *Rosa26* (Figure S1A). RNA-Seq after treatment with 1000ng/mL dox demonstrated that in both clones, *Airn* expression was induced to levels that slightly surpassed its endogenous levels in trophoblast stem cells (TSCs; Figures 1B, S1B). Like endogenous *Airn* in TSCs ^4^, ectopically expressed *Airn* localized as a single focus at its presumptive site of transcription, underwent transcriptional attrition across its gene body, rarely underwent splicing, and was unstable, with a half-life of ∼45 minutes (Figures 1C-E, 1G). We conclude that ectopically expressed *Airn* retains properties that are similar to the endogenous transcript.

**Figure 1.**
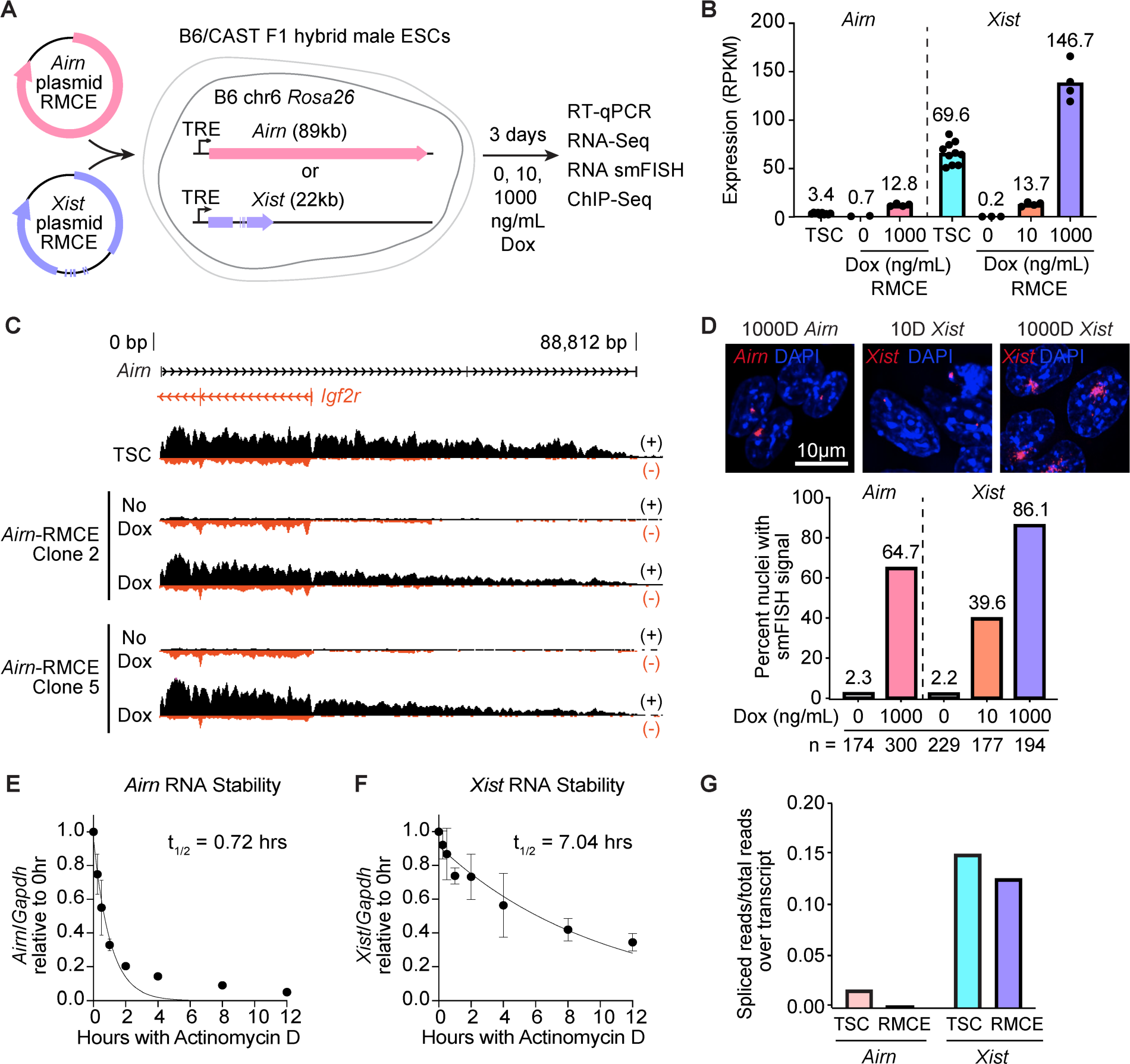
Ectopically expressed *Airn* resembles the endogenous *Airn* transcript. **(A)** Schematic of *Airn* and *Xist* gene insertion into the *Rosa26* locus on B6 chr6 in F1-hybrid B6/CAST ESCs via RMCE. **(B)** RNA-Seq RPKM expression levels of *Airn* and *Xist* in TSCs and in RMCE ESCs following treatment with various concentrations of dox. RPKM, Reads Per Kilobase of transcript, per Million mapped reads. **(C)** Genome browser scaled view of strand-specific RNA-Seq signal from *Airn* in TSCs versus untreated and dox-treated *Airn*-RMCE ESC clones. **(D)** Representative *Xist* and *Airn* RNA smFISH images (top) and counts of nuclei with detectable smFISH signal (bottom) for RMCE ESCs treated with the indicated concentration of dox, labeled with Quasar 570-conjugated probes (red) and DAPI (blue). Scale, 10μm. **(E-F)** Half-life measurements for *Airn* and *Xist* RNA in RMCE ESCs determined by RT-qPCR following treatment with actinomycin D for the indicated time. **(G)** Extent of splicing of *Xist* and *Airn* transcripts in TSCs and RMCE ESCs.

We next compared the relative expression levels of ectopically expressed *Airn* to those of *Xist*. We had previously used the same Cre-mediated approach to insert the 22kb *Xist* gene into *Rosa26* under control of the same dox-inducible promoter used to express *Airn* (Figure S1A) ^12^. In these ESCs, after treatment with 1000ng/mL dox, we found that *Xist* was expressed at levels ∼10-fold higher than *Airn*, consistent with its ∼10-fold greater half-life (Figures 1B, 1E, 1F). After treatment with only 10ng/mL dox, *Xist* was expressed at levels on par with *Airn* expressed with 1000ng/mL dox (Figure 1B). RNA FISH demonstrated that at the 1000ng/mL condition, 65% and 86% of cells expressed detectable levels of *Airn* and *Xist*, respectively (Figure 1D). At the 10ng/mL condition, 40% of cells expressed detectable levels of *Xist* (Figure 1D). Expression levels of the reverse tetracycline-controlled transactivator (rtTA) were equivalent across genotypes, supporting the notion that each *Airn* and *Xist* ESC line was capable of inducing lncRNA expression to equivalent extents (Figure S1C). We conclude that when inserted into the same genomic locus, under control of the same promoter, and subject to the same level of promoter induction, ectopically expressed *Airn* is less stable and accumulates to levels lower than *Xist*.

We next performed ChIP-Seq to determine the extent to which ectopic expression of *Airn* altered surrounding levels of PRC2-deposited H3K27me3. In addition to examining H3K27me3 levels in *Airn*-expressing cells, we examined H3K27me3 levels in “Empty” control ESCs harboring a dox-inducible promoter at *Rosa26* but no inserted lncRNA, as well as in *Xist*-expressing cells treated without dox or with 10ng/mL or 1000ng/mL dox (Figure 2A). Relative to Empty control ESCs, in *Airn*-expressing ESCs, we observed a striking increase of H3K27me3 over the majority of chr6, specifically on the B6 and not CAST allele (Figures 2A, 2B, S2A). Moreover, the increases in H3K27me3 induced by *Airn* were significantly larger than those induced by *Xist* when the two lncRNAs were expressed at near-equivalent levels (1000ng/mL dox for *Airn*, 10ng/mL dox for *Xist*; Figures 1B, 2A, 2B). The increases in H3K27me3 when *Xist* was expressed at 1000ng/mL dox surpassed those of *Airn* at the 1000ng/mL condition (Figures 2A, 2B). We conclude that the *Airn* lncRNA product is functional, its expression being sufficient to induce the deposition of H3K27me3 over the majority of chr6 when expressed from *Rosa26*. Moreover, in this context and when normalized for copy number, *Airn* deposits H3K27me3 with a potency that rivals or even surpasses that of *Xist*.

**Figure 2.**
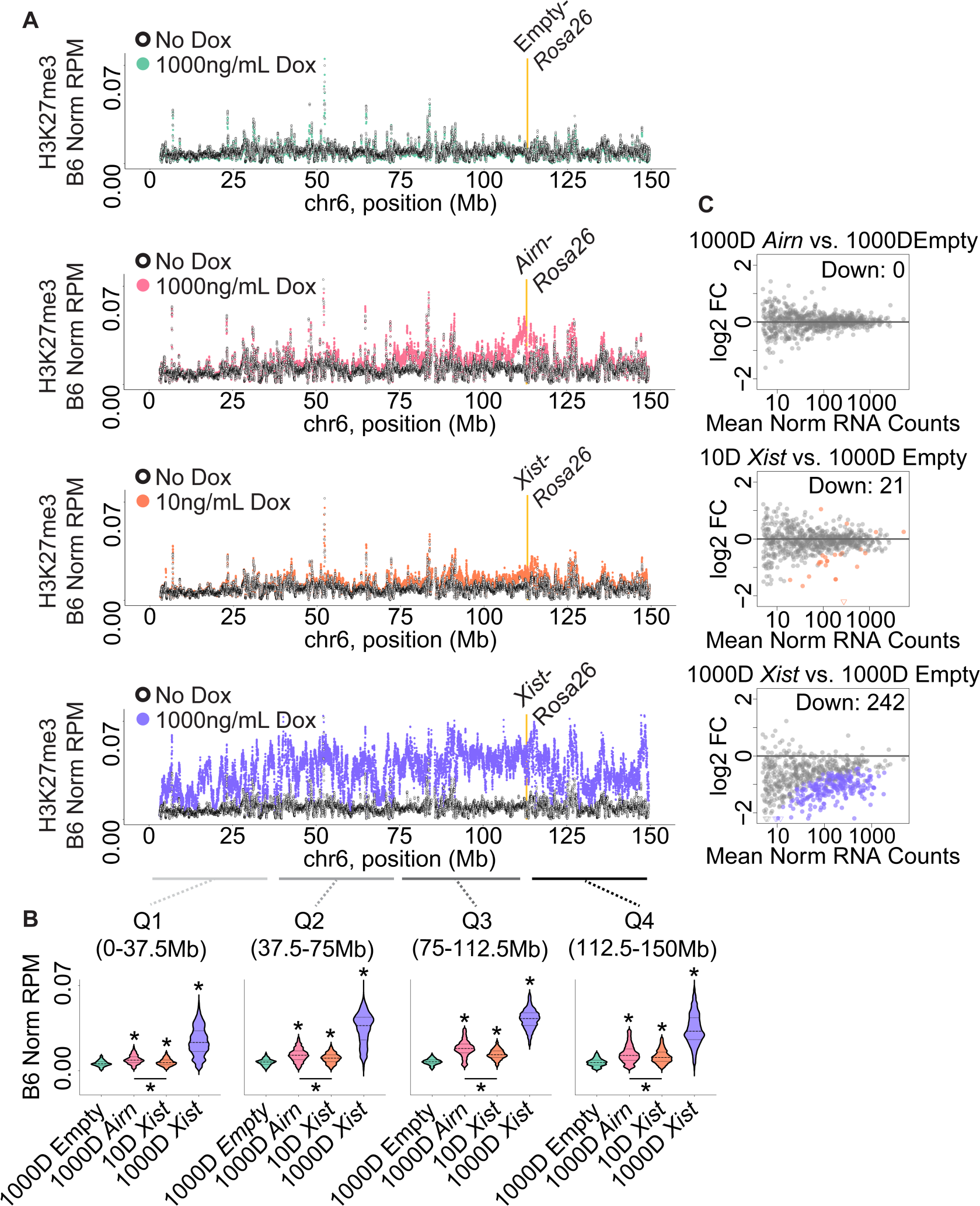
Changes in chr6 H3K27me3 density and gene expression induced by *Airn* and *Xist*. **(A)** Tiling density plots of H3K27me3 ChIP-Seq signal over B6 chr6 in untreated and dox-treated RMCE ESCs. Yellow bar, *Rosa26* locus: RMCE cargo insertion site. **(B)** H3K27me3 levels on B6 chr6 in dox-treated RMCE ESCs in binned quartiles. Asterisks, TukeyHSD adjusted p value < 0.0001; top asterisks represent comparisons to 1000D Empty. **(C)** MA plots showing differential expression of B6 chr6 genes relative to 1000ng/mL dox-treated empty-cargo RMCE ESCs. Log fold change was plotted against the mean of normalized B6 counts. Each dot is an expressed gene, and colored dots show those that are significantly changed as determined by DESeq2 analysis (adjusted p value < 0.01).

We next examined our RNA-Seq data to determine the extent to which *Airn* and *Xist* repressed gene expression. Despite the large-scale changes in H3K27me3, *Airn* did not induce repression of any chr6 genes (Figure 2C, Table S1). Likewise, prolonged, 7-day *Airn* expression did not induce repression (Table S1). In contrast, at 10ng/mL dox, *Xist* repressed expression of 21 chr6 genes, and at 1000ng/mL dox, *Xist* repressed expression of 242 chr6 genes (Figure 2C, Table S1). Thus, *Airn*’s ability to repress transcription is markedly weaker than that of *Xist*. Also, *Xist*’s ability to repress transcription correlates positively with its abundance.

Our study unequivocally demonstrates that the *Airn* lncRNA product is functional, being capable of inducing the accumulation of Polycomb over a genomic range and intensity that rivals that of *Xist*. This is despite the fact that *Airn* and *Xist* harbor diverse evolutionary origins and sequence contents ^13^. We also demonstrate that under the conditions tested above, *Airn* does not induce transcriptional repression despite its ability to induce deposition of H3K27me3. Transcriptional repression and H3K27me3 deposition have been shown to be separable functions within *Xist* ^14–17^. Our observation that the same is true for regulation by *Airn* suggests that these principles are generalizable to other lncRNAs, and supports the possibility that in its endogenous locus, *Airn*, along with the Polycomb-directed marks that it helps to deposit, functions largely to maintain a memory of previous transcriptionally repressive events ^18, 19^. We speculate that the establishment of transcriptional memory may be a common function of *cis*-acting lncRNAs, perhaps more common than the type of robust, *de novo* gene silencing induced by *Xist*.

Lastly, our data underscore the importance of RNA abundance in moderating the extent of lncRNA-mediated repression ^4^. In the context of *Rosa26,* when subject to identical extents of promoter activation, *Xist* accumulated to levels that were 10-fold higher than those of *Airn*. These elevated levels were necessary for *Xist*’s full repressive effect. We speculate that the evolution of full-scale X inactivation ^20^ may have been driven, in part, by the acquisition of features within *Xist* that led to punctuated increases in its nuclear abundance. Conversely, without such features, lncRNAs such as *Airn* effectively limit their influence on chromatin.

## ACKNOWLEDGEMENTS

This work was supported by NIH grant R01GM136819 and RO1GM121806 (J.M.C.). J.B.T. was supported by T32CA217824. A.K.B was supported by T32GM119999 and F31HD103370. R.E.C. was supported by T32GM007092 and F31HD103334. M.M.M. was supported by F31HD111292 and T32GM119999. S.B.P. and E.W.A. were supported by T32GM135128.

## AUTHOR CONTRIBUTIONS

J.B.T., A.K.B. and J.M.C. conceived the study; J.B.T., A.K.B., S.R.B., S.P.B., R.E.C., M.M.M., Q.E.E., E.W.A., D.O.C., and J.M.C. performed the experiments; D.O.C. provided reagents and advice; and J.B.T., A.K.B., and J.M.C. wrote the paper with feedback from co-authors.

## Figures and legends

**Figure S1.**
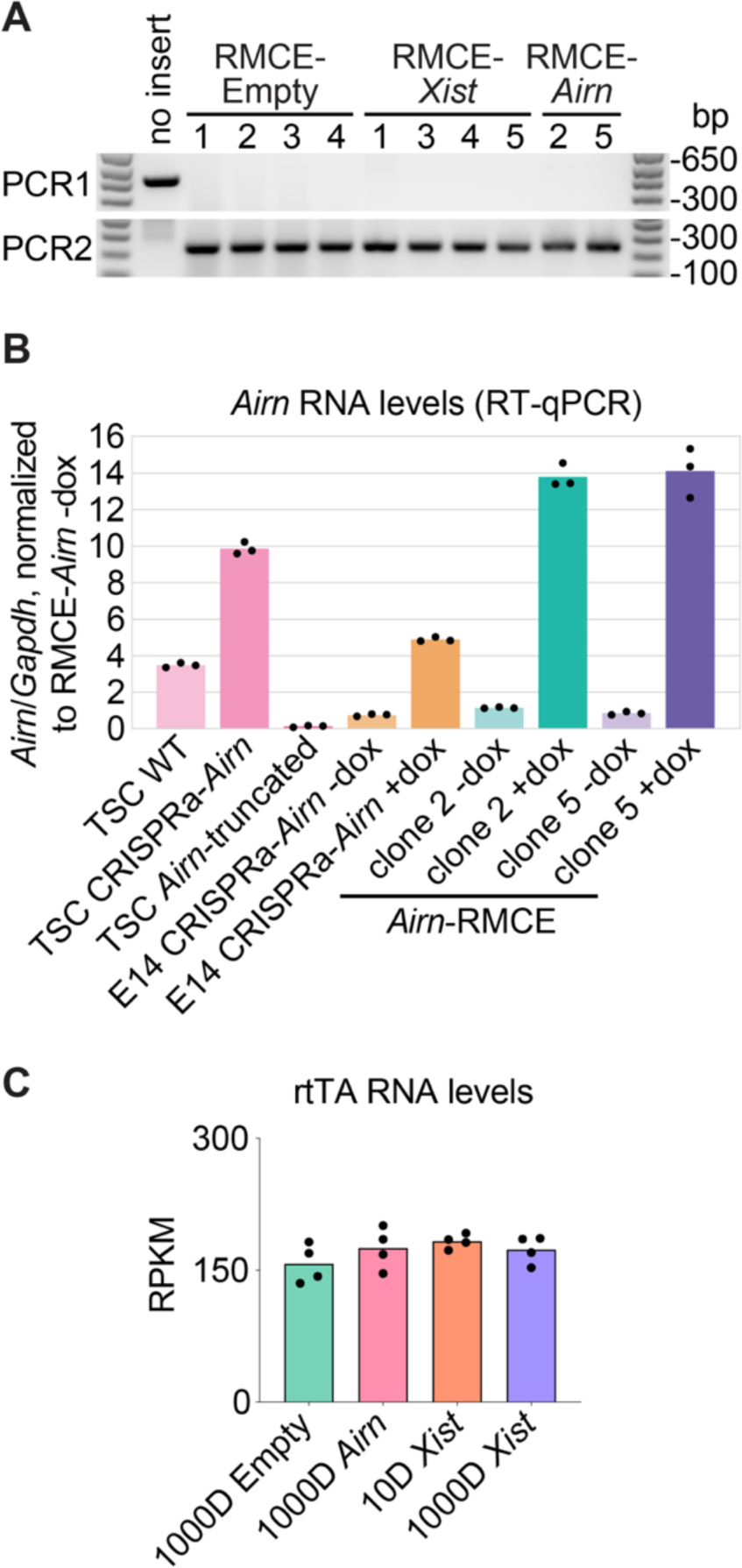
**(A)** Genotyping PCR products confirming successful insertion of cargo sequences into B6 *Rosa26* via RMCE. Lanes are labeled with numbers corresponding to the ID of each clonal cell line. **(B)** RT-qPCR comparison of *Airn* levels in RMCE ESCs to those in TSC and ESC cell lines from previous studies ^4, 21^. **(C)** RNA-Seq RPKM expression levels of rtTA across dox-treated RMCE ESCs.

**Figure S2.**
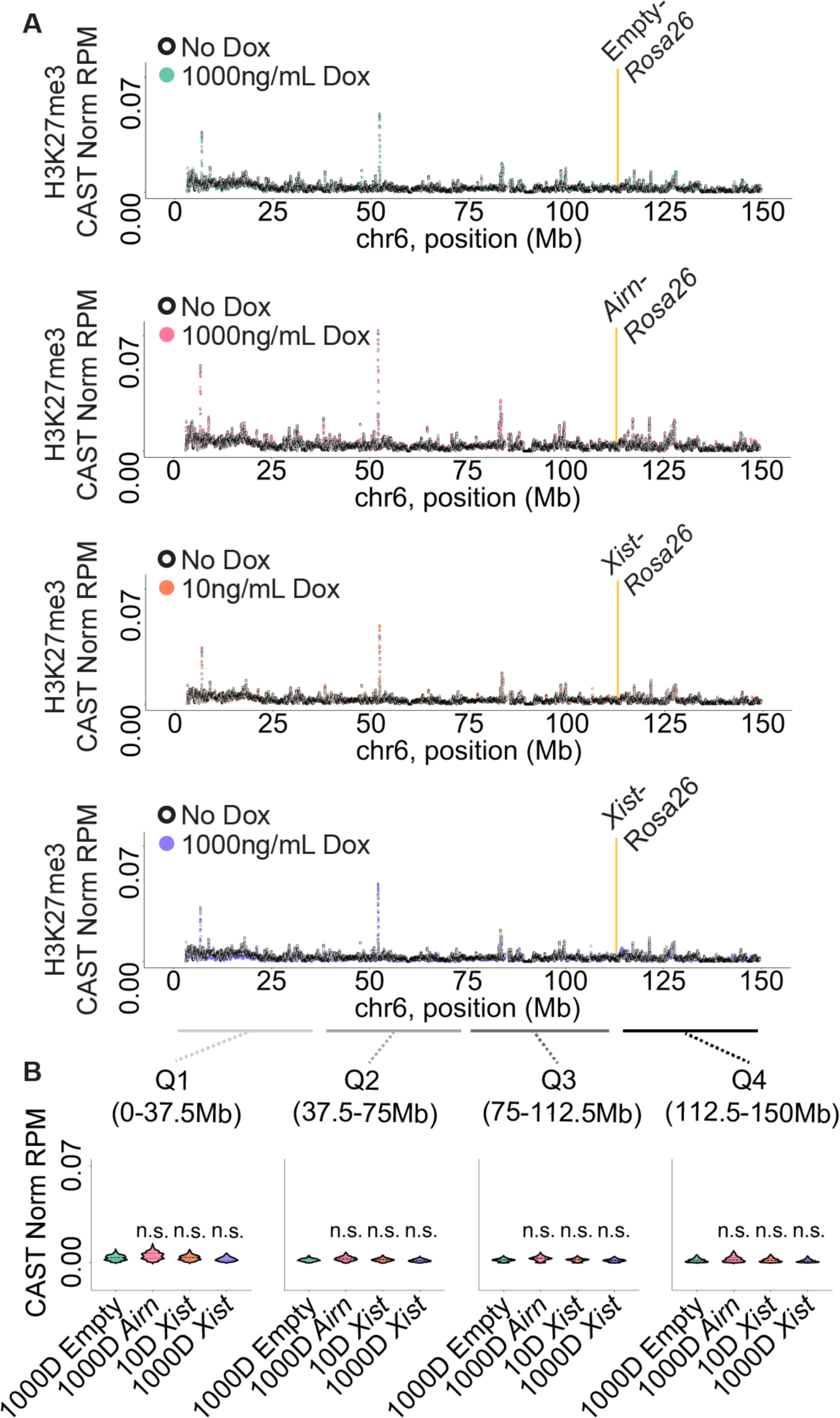
**(A)** Tiling density plots of H3K27me3 ChIP-Seq signal over CAST chr6 in untreated and dox-treated RMCE ESCs. Yellow bar, *Rosa26* locus: RMCE cargo insertion site. **(B)** H3K27me3 levels in CAST chr6 across dox-treated RMCE ESCs in binned quartiles. Asterisks, TukeyHSD adjusted p value < 0.0001, as in Figure 2B; top asterisks represent comparisons to 1000D Empty.

## Methods

### pCARGO-RMCE-Airn Plasmid construction

BAC clone RP23-309H20 harboring the Airn lncRNA gene was obtained from the BACPAC Resources Center and was modified by recombineering to generate the tetracycline-inducible pCARGO-RMCE-*Airn* donor vector. In brief: a first recombineering step was performed to remove the lox511 site in the pBACe3.6 vector backbone along with the ∼56 kb of mouse genomic sequence located upstream of *Airn*/ NR_027772 exon 1 (mm10 chr17:12741311). The removed sequences were replaced by an ampicillin resistance cassette, a lox71 site, and a TreTight/CMV minimal promoter, all from the pCARGO-RMCE cargo vector described in ^12^, immediately upstream of *Airn* exon 1. A second recombineering step was performed to remove ∼119 kb of mouse sequence downstream of the transcription end site for *Airn*/ NR_027772 (mm10 chr17:12830123) as well as the loxP site in the BAC vector backbone. The removed sequences were replaced by an SV40 polyadenylation sequence, an FRT-flanked PGK-EM7-Hygromycin resistance cassette, and a lox2272 site, again from the pCARGO-RMCE cargo vector described in ^12^. The final BAC clone was fully sequence-validated. Large-scale preparation of pCARGO-RMCE-*Airn* plasmid was purified from cells grown overnight at 30°C with ampicillin selection (100μg/mL ampicillin sodium salt, Fisher Scientific BP1760), using the NucleoBond BAC100 Maxiprep Kit (Macherey-Nagel 740579). Pelleted plasmids were resuspended in Invitrogen TE Buffer before measuring DNA concentration with a Qubit 2.0 Fluorometer (dsDNA high sensitivity kit, Thermo Fisher Q32851). Integrity and sequence composition of plasmid preparations were confirmed by diagnostic digestion and by whole-plasmid Oxford Nanopore sequencing (Plasmidsaurus, File S1). Full details of the BAC recombineering approach are provided in File S2.

### Cell culture

B6/CAST F1-hybrid *Rosa26*-RMCE ESCs from ^12^ were grown on gelatin-coated plastic dishes in a humidified Thermo Fisher Forma Series II water-jacketed incubator at 37°C and under 5% CO_2_. Cells were grown in DMEM high glucose plus sodium pyruvate (Gibco 11995-065), 15% ESC-qualified fetal bovine serum (Gibco 26140-079), 0.1mM non-essential amino acids (Gibco 11140-050), 100U/mL penicillin-streptomycin (Gibco 15140-122), 2mM L-glutamine (Gibco 25030-081), 0.1mM β-mercaptoethanol (Sigma-Aldrich 63689), and 1:500 LIF conditioned media produced from Lif-1Cα (COS) cells.

TSCs were cultured as in (Quinn et al. 2006). Briefly, TSCs were cultured on gelatin-coated, pre-plated irradiated mouse embryonic fibroblast (irMEF) feeder cells in TSC media (RPMI [Gibco 11875093], 20% qualified FBS [Gibco 26140079], 0.1mM penicillin-streptomycin [Gibco 15140122], 1mM sodium pyruvate [Gibco 11360070], 2mM L-glutamine [Gibco 25030081], 100μM β-mercaptoethanol [Sigma-Aldrich 63689]) supplemented with 25ng/mL FGF4 (Gibco PHG0154) and 1μg/mL Heparin (Sigma-Aldrich H3149) just before use, at 37°C in a humidified incubator at 5% CO_2_. At passage, TSCs were trypsinized with 0.125% Trypsin-EDTA in PBS solution (Gibco 25200-072) for ∼4 minutes at room temperature and gently dislodged from the plate with a sterile, cotton-plugged Pasteur pipette. To deplete irMEFs from TSCs prior to all harvests, TSCs were pre-plated for 45 minutes at 37°C, transferred to a fresh culture plate, and then cultured for three days in 70% irMEF-conditioned TSC media supplemented with growth factors as above.

### Generation of RMCE cell lines

Prior to electroporation, plasmid mixtures were prepared as follows: for RMCE-empty cells: 6.25μg pCARGO-RMCE-empty ^12^, 4.1μg pOG-Cre, 6μg PB-TRE-Cas9 (for transient hygromycin B resistance ^22^; for RMCE-*Xist* cells: 19μg pCARGO-RMCE-*Xist* ^12^, 2.2μg pOG-Cre, 6μg PB-TRE-Cas9; for RMCE-*Airn* cells, two conditions that each ultimately yielded one colony: (19μg pCARGO-RMCE-*Airn*, 2.2μg pOG-Cre, 6μg pCL116 ^22^) or (54 μg pCARGO-RMCE-*Airn*, 10.8μg pOG-Cre, 6μg pCL116). Each plasmid mixture was ethanol-precipitated, air-dried, and thoroughly resuspended in 10μL of TE Buffer (Invitrogen) after soaking overnight at 4°C. The 10-μL plasmid mixtures were mixed with 1 million B6/CAST F1-hybrid *Rosa26*-RMCE ESCs (in 100 μL Neon Buffer R) and electroporated using a Neon Transfection System (Invitrogen) with one 40-ms pulse of 1000 V before seeding sparsely onto a confluent 10-cm dish of gamma-irradiated drug-resistant (DR4) MEF feeder cells (ATCC SCRC-1045) in growth medium lacking penicillin-streptomycin. Growth medium was not changed the day after electroporation but was replaced daily thereafter. Two days after electroporation, growth medium was replaced with normal growth medium (containing penicillin-streptomycin). Starting three days after electroporation and lasting for one day (RMCE-*Airn*) or at least ten days (RMCE-Empty and RMCE-*Xist*), cells were selected in growth medium containing 150μg/mL hygromycin B (Roche 10843555001). Ganciclovir (InvivoGen sud-gcv) was added at a working concentration of 3μM starting five days (RMCE-*Airn* cells electroporated with 54μg pCARGO-RMCE-*Airn* plasmid), six days (RMCE-*Airn* cells electroporated with 19μg pCARGO-RMCE-*Airn* plasmid), or seven days (RMCE-Empty and RMCE-*Xist*) after electroporation and lasting for least six days. When clearly visible by eye (9-12 days after electroporation), individual colonies were picked with aid of a light microscope fitted with a 4X objective, dissociated with 0.125% trypsin, and each transferred to a 24-well plate well pre-seeded confluently with gamma-irradiated DR4 MEF feeder cells. Clonal lines of cells were grown until sufficiently confluent (>50%), at which point they were passaged for subsequent genotyping, preparing cryogenic freeze-down stocks, and introducing the reverse tetracycline-controlled transactivator (rtTA) via plasmid transfection.

For introduction of rtTA, four independent RMCE-Empty cell lines, four independent RMCE-*Xist* wild-type cell lines, and two independent RMCE-*Airn* cell lines were each seeded at a density of 750,000 cells per well of a 6-well plate. The following day, each cell line was given fresh growth medium and transfected with a plasmid mixture containing 1μg PB-rtTA (Addgene #126034; ^22^) and 1μg pUC19-piggyBac transposase ^23^ using Lipofectamine 3000 (Thermo Fisher L3000001). Starting around 24h after transfection and lasting at least 10 days, cells were selected in growth medium containing 200μg/mL G418 sulfate (Thermo Fisher 10131035).

### Genomic DNA preparation and genotyping PCR

To prepare genomic DNA, cells in a confluent well of 24-well plate were lysed by adding 488μL of a mixture containing 400μL lysis buffer (100 mM Tris-HCl pH 8.0, 5mM EDTA, 200mM NaCl, 0.2% (w/v) SDS), 80μL 20mg/mL proteinase K (Meridian Bioscience BIO-37084), and 8μL 5mg/mL linear acrylamide (Thermo Fisher AM9520) and incubating at 55 °C overnight. Samples were then heated at 100 °C for 1h, incubated at 4 °C for at least 30min, and centrifuged at 16,100 x g for 5min at 4°C. Supernatants were removed, and pellets were washed with 1mL ice-cold 80% ethanol (v/v). After centrifuging again as before, supernatants were completely removed, and pellets were air-dried before adding 100μL TE Buffer (Thermo Fisher 8019005) and incubating at 4°C overnight. Genomic DNA samples were mixed thoroughly by vortexing and pipetting before using for genotyping PCR.

To perform genotyping PCR, 25-μL reactions were prepared on ice in PCR strip-tubes, each containing the following: 1μL 10mM dNTPs, 0.25μL forward primer, 0.25μL reverse primer (see Table S3 for oligonucleotide sequences), 2.5μL 10X Ammonium Buffer, 0.75μL 50 mM MgCl2, 18.25 nuclease-free water, 0.5μL 1U/μL Apex Taq Polymerase (Genesee Scientific 42-401), and 1.5μL genomic DNA. Reactions were run on a thermocycler with the following program: 95 °C for 3min, 30 cycles of (95°C for 30s, 60°C for 30s, 72°C for 1min), 72°C for 5min. To each PCR product, 5μL 6X gel loading dye (NEB B7024S) was added, and a 10-μL sample of each was run on a 1.5% agarose-TAE-ethidium gel alongside 5μL Invitrogen 1 Kb Plus DNA Ladder (Thermo Fisher 10787018) before imaging with a Bio-Rad ChemiDoc MP Imaging System (Ethidium Bromide mode, 0.1s exposure).

### RNA preparation

Prior to harvesting, RMCE cell lines (with rtTA) were treated with 0-1000ng/mL doxycycline (1000ng/mL if not specified otherwise) for 72 hours (if not specified otherwise). At harvest, RMCE cells or MEF feeder-free TSCs (see above) were washed twice with ice-cold PBS and harvested with 1 mL TRIzol (Thermo Fisher 15596026). Total RNA was prepared via standard TRIzol-chlorofom extraction, per manufacturer’s instructions, except for the addition of 4μL 5μg/μL linear acrylamide (Thermo Fisher AM9520) to promote precipitation in isopropanol. RNA pellets were fully resuspended in RNase-free water, RNA concentrations were measured via Qubit 2.0 fluorometer (RNA high sensitivity kit, Thermo Fisher Q32852), and integrity of RNA was confirmed by visualizing rRNA bands on an agarose gel. Working stocks of 100ng/μL total RNA were prepared, and all RNA samples were stored at -80°C.

### Reverse transcription and quantitative PCR (RT-qPCR)

Using the Applied Biosystems High-Capacity cDNA Reverse Transcription Kit (Thermo Fisher 4368814), 20-μL randomer-primed reverse transcription reactions were prepared in 0.2-mL PCR strip-tubes on ice, each containing 10μL 100ng/μL total RNA (1000 ng) and 0.5μL 40U/μL RNaseOUT RNase Inhibitor (Thermo Fisher 10777019). Reverse transcription was performed with the following thermal cycler program: 25°C for 10min, 37°C for 120min, 85°C for 5min, hold at 4°C. For robust quantification via qPCR, a standard curve was constructed by pooling a portion of representative reverse transcription product samples and serially diluting this mixture 4-fold with nuclease-free water to generate 6 standards. For quantification of each sample, reverse transcription products were each diluted 10-fold with nuclease-free water. For each qPCR primer pair, a qPCR master mix was generated, containing for each 10-μL qPCR reaction: 5μL iTaq Universal SYBR Green Supermix (Bio-Rad 1725124), 3μL nuclease-free water, 0.5μL 10μM forward primer, and 0.5μL 10μM reverse primer (see Table S3 for oligonucleotide sequences) ^4, 12^. In Hard-Shell 96-Well PCR Plates (Bio-Rad HSP9601), qPCR reactions were set up in technical triplicate by combining 9μL qPCR master mix and 1μL of standard-curve or 10-fold-diluted reverse transcription product. A standard curve was run for each primer pair on every plate. Plates were firmly sealed with Microseal ’B’ PCR Plate Sealing Film (Bio-Rad MSB1001), mixed several times by inversion and low-intensity vortexing, and briefly pulsed down in a centrifuge (up to 2000 x g) to collect the volumes and remove bubbles. Reactions were analyzed on a Bio-Rad C1000 Touch Thermal Cycler equipped with a CFX96 Real-Time System with the following program: 95°C for 10min, 40 cycles of (95°C for 15s, 60°C for 30s, 72°C for 30s, plateread), followed by melt curve analysis: 95°C for 10s, 65°C for 31s, 61 cycles of (65°C for 5s +0.5°C/cycle Ramp 0.5°C/s, plateread). Data were analyzed using Bio-Rad CFX manager, first setting “Cq Determination Mode” to “Regression” and then using each primer pair’s standard curve (Cq vs. starting quantity) to convert Cq values to starting quantities for each technical replicate. Average *Airn* starting quantities were normalized by dividing by the average *Gapdh* starting quantity for each biological sample, and *Gapdh*-normalized *Airn* expression data were plotted using GraphPad Prism.

### Single-molecule RNA FISH

Stellaris RNA FISH probes were designed against the first 40kb of *Airn* utilizing the Stellaris RNA FISH Probe Designer (LGC, Biosearch Technologies, Petaluma, CA) available online at www.biosearchtech.com/stellarisdesigner (version 4.2; parameters: masking level 5; max number of 48 probes; oligo length 20nt; min. spacing length 2nt; Quasar 570). Probes to *Xist* were designed by Stellaris (product # SMF-3011-1, Quasar 570). ESCs were hybridized with the Stellaris RNA FISH probes following the manufacturer’s instructions for adherent cells, available online at www.biosearchtech.com/stellarisprotocols. Briefly, cells were grown on coverslips, fixed with 3.7% formaldehyde and permeabilized overnight in 70% ethanol. Cells were then incubated in Wash Buffer A with formamide for five minutes at room temperature, before hybridization with 125nM of *Airn* or *Xist* probes in hybridization buffer at 37°C for 4 hours, then washed twice for 30 minutes at 37°C with Wash Buffer A. Cells were then washed for 5 minutes at room temperature once with Wash Buffer B, then fixed with ProLong Gold containing DAPI (Thermo Fisher P36941) overnight at room temperature in the dark.

Two biological replicates per doxycycline condition were imaged. Images were taken with a Leica DMi8 inverted confocal microscope, using Leica Application Suite X software version 3.7.5.24914. Images used for counting *Xist* or *Airn* expression in nuclei were taken with 40x or 63x objectives. Representative images were taken with 100x objective. Z-slice sizes were 0.2 µm. Images were deconvolved with Huygens Essential version 20.04.0p3 64b (Scientific Volume Imaging, The Netherlands, http://svi.nl) using the standard deconvolution profile under batch express. Images were analyzed using FIJI ^24^. To quantify nuclei with or without *Xist* or *Airn* expression, the 570 channel was turned off, DAPI channel was adjusted so that distinct nuclei were visible and then numbered. Then DAPI was turned off, and the 570 channel was adjusted so that signal inside nuclei, but not background signal was visible. Numbered nuclei were then counted as having *Xist*/*Airn* signal or not.

### RNA half-life measurement

RNA half-lives were measured using Actinomycin D. *Airn*-RMCE and *Xist*-RMCE cells were induced for 2 days with 1000ng/mL doxycycline prior to start of the Actinomycin D time course. ESCs were then treated with 5µg/mL Actinomycin D (Sigma-Aldrich A9415-5MG) for 15 minutes, 30 minutes, 1 hour, 2 hours, 4 hours, 8 hours, and 12 hours (or without Actinomycin D, “0hr”) and lysed with TRIzol. RNA was extracted and subjected to RT-qPCR (see above) to measure levels of *Airn*, *Xist*, and *Gapdh* at each time point (see Table S3 for oligonucleotide sequences). Time course experiments were done in biological replicate for both cell types.

RNA levels were normalized to *Gapdh* at each time point and calculated as the percentage of RNA relative to the 0hr time point. To estimate half-life, *Gapdh*-normalized data for each replicate were averaged and fit to a non-linear one-phase decay model in GraphPad Prism using the equation Y=(Y0)*exp(-K*X). Error bars represent the standard deviation between biological replicates.

### RNA-Seq

Libraries for RNA sequencing were prepared using the KAPA RNA HyperPrep Kit with RiboErase (HMR) (Roche 08098140702) and KAPA Unique Dual-Indexed Adapters (Roche 08861919702) or TruSeq adapters, with each library’s input consisting of 9μL 100ng/μL total RNA (900 ng) mixed with 1μL of a 1:250 dilution of ERCC RNA spike-in controls (Thermo Fisher 4456740). RNA fragmentation was performed at 94°C for 6min, and libraries were amplified using 11 cycles of PCR. Libraries were quantified using a Qubit 2.0 fluorometer (dsDNA broad sensitivity kit, Thermo Fisher Q32850), pooled, and sequenced on an Illumina NextSeq500 platform with a 75-cycle NextSeq 500/550 High Output Kit v2.5 (Illumina 20024906).

Sequencing reads (fastq files) were mapped both to the GENCODE vM25 mouse (mm10) genome (“basic CHR” from https://www.gencodegenes.org/mouse/release_M25.html) [for B6] and to a version of the same genome modified to incorporate CAST single-nucleotide polymorphisms (SNPs) [for CAST]. Mapping was performed with STAR (v2.7.9a ^25^), using two-pass mapping and the option “--outFilterMultimapNmax 1” to consider only uniquely mapping reads. Using a custom Perl script (intersect_reads_snps16.pl) and a file containing a list of CAST SNPs (downloaded from the Sanger Mouse Genomes Project on 7/30/2020 ^26^), aligned reads in the B6 and CAST SAM files were parsed to identify reads clearly originating from either the B6 or CAST allele (i.e. reads that overlap at least 1 B6/CAST SNP). Reads marked as either B6 or CAST were then assigned to genes using a custom Perl script (ase_analyzer10.pl) and the GTF file gencode.vM25.basic.annotation.gtf ^27^. The number of B6-specific reads for each gene from each sample were then compiled. Prior to running DESeq2, a pre-filtering step was used in which genes with fewer than an average of five B6 reads per sample were excluded. Using R, DESeq2 ^28^ was used with default settings, except for changing the significance threshold to an adjusted p value of 0.01, to determine differential gene expression and to generate MA plots. For RPKM expression analysis, total, non-allelic reads were counted over gene annotations from gencode.vM25.annotation.gtf using featureCounts with default parameters ^29^. Read counts for each gene were then divided by total number of reads in the dataset, divided by a million, and then divided by the gene length.

### Quantification of spliced reads

Aligned reads with quality score over thirty were merged from all replicates. Samtools ^30^ was used to extract reads over *Airn* and *Xist*, and count the reads over each transcript. Reads with gap junctions greater than or equal to 125 bp were then extracted and labeled as spliced reads. The number of spliced reads over *Airn* and *Xist*, were then divided by total merged reads over each transcript, respectively.

### H3K27me3 ChIP-Seq

Prior to crosslinking, RMCE cell lines (with rtTA) were treated with 0-1000ng/mL doxycycline (1000ng/mL if not otherwise specified) for 72 hours (or longer, 7 days). Adhered cells were crosslinked with 0.6% formaldehyde (Fisher Scientific BP531-500) in DMEM media with 5% FBS for 10min at room temperature, then quenched with 125mM glycine for 5min at room temperature. Crosslinked cells were then washed twice with ice-cold PBS and scraped with ice-cold PBS with 0.05% Tween (Fisher Scientific EW-88065-31) and PIC (Sigma Aldrich P8340). The cells were then spun at 1,200 x g at 4°C to remove PBS, followed by resuspension in ice-cold PBS with PIC and divided into 10-million cell aliquots. Each ChIP was performed using 10 million cells, 10μL H3K27me3 antibody (Cell Signaling 9733), and 30μL Protein A/G agarose beads (Santa Cruz sc-2003). Antibody-conjugated beads were prepared by incubating antibody with beads in 300μL Blocking Buffer (PBS, 0.5% BSA [Invitrogen AM2616]) overnight at 4°C with rotation.

10 million crosslinked cells were resuspended in 1mL Lysis Buffer 1 (50mM HEPES pH 7.5, 140mM NaCl, 1mM EDTA, 10% glycerol, 0.5% NP-40, 0.25% Triton X-100, PIC) and incubated with rotation for 10min at 4°C. Cells were then resuspended in 1mL Lysis Buffer 2 (10mM Tris-HCl pH 8.0, 200mM NaCl, 1mM EDTA, 0.5 mM EGTA, PIC) for 10 minutes at room temperature. All buffer removal steps were performed with 5-min 1,200 x g spins at 4°C. The extracted nuclei pellet was then resuspended in 500μL Lysis Buffer 3 (10mM Tris-HCl pH 8.0, 100mM NaCl, 1mM EDTA, 0.5mM EGTA, 0.1% sodium-deoxycholate, 0.5% N-lauroylsarcosine, PIC), and chromatin was sonicated to 100-500bp fragments using a Vibracell VX130 (Sonics) with the following parameters: 10 cycles of 30% intensity for 30 seconds with 1 minute of rest on ice between cycles. Soluble chromatin was obtained with a 30-min max speed spin at 4°C, mixed with 1% Triton 0, and then incubated with pre-conjugated antibody beads overnight at 4°C with rotation. The ChIP beads were washed five times in 1mL RIPA Buffer (50mM HEPES pH 7.5, 500mM LiCl, 1mM EDTA, 1% NP-40, 0.7% sodium-deoxycholate, PIC) and once with 1mL TE, each for 5 minutes at 4°C with rotation and spun at 2,000 x g for 2 minutes for buffer removal. The washed beads were then resuspended in Elution buffer (50mM Tris-HCl pH 8.0, 10mM EDTA, 1% SDS) and placed on a 65°C heat block for 17 minutes with frequent vortexing. ChIP DNA was then reverse crosslinked in 0.5% SDS and 100mM NaCl overnight at 65°C, followed by a 1-h RNaseA (3μL; Thermo Scientific EN0531) treatment at 37°C and a 2.5-h Proteinase K (10μL; Invitrogen 25530015) treatment at 56°C. DNA was then extracted with 1 volume of phenol:chloroform:isoamyl alcohol (Sigma-Aldrich P3803) and precipitated with 2 volumes 100% ethanol, 1/10 volume 3M sodium-acetate pH 5.4, and 1/1000 volume linear acrylamide (Invitrogen AM9520) overnight at -20°C. Precipitated DNA was then extracted with a 30-minute max speed spin at 4°C, washed once with ice-cold 80% ethanol, and resuspended in TE.

ChIP-Seq libraries were prepared with NEBNext End Repair Module (NEB E6050S), A-tailing by Klenow Fragment (3’è5’ exo-; NEB M0212S), and TruSeq 6-bp index adaptor ligation by Quick ligase (NEB M2200S), and NEBNext High-Fidelity 2X PCR Master Mix (NEB M0541S). All DNA clean-up steps were performed with AMPure XP beads (Beckman Coulter A63880). Single-end, 75-bp sequencing was performed using an Illumina NextSeq 500/550 High Output v2.5 kit (Illumina 20024906) on a NextSeq 500 System.

ChIP-Seq reads were aligned using bowtie2 with default parameters ^31^. Aligned reads that had a mapping quality greater than or equal to 30 were extracted with samtools ^30^, and allele-specific read retention (i.e., reads that overlap at least one B6 or CAST SNP) was performed as in ^32, 33^ using a custom Perl script (intersect_reads_snps16.pl). For chromosome-scale tiling density plots, B6- or CAST-specific reads were summed in 10kb bins across chr6. Binned counts were then divided by the total number of reads in the dataset and divided by a million (i.e., RPM), then divided by the number of B6/CAST SNPs detected in the bin genomic coordinates (i.e., SNP-norm RPM). Finally, bins were averaged every 9 bins in 1bin increments. For plotting, only bins with greater or equal to 25 B6/CAST SNPs were retained for higher confidence interpretation of the allelic data. All plots were generated using ggplot2 ^34^ in RStudio. For statistical comparisons of H3K27me3 levels across conditions, SNP-norm RPM signal of chr6 bins were divided by quartiles, then subset for every 10^th^ bin. An ANOVA-TukeyHSD test was applied to all B6 and CAST data for adjusted p values; two conditions were considered significantly different with an adjusted p value threshold of < 0.0001.

## DATA ACCESSION

Data are deposited under GEO accession GSE230442.

